# Decreased female survival may explain wild turkey decline

**DOI:** 10.1101/2025.05.16.654534

**Authors:** Marcus A. Lashley, M. Colter Chitwood, Anna K. Moeller, William D. Gulsby, Alex D. Potash, Kelly O’Neil, Mark Turner

## Abstract

Recent declines in wild turkey (*Meleagris gallopavo*) populations have prompted extensive research efforts and adjustments to state hunting regulations across the range of wild turkeys. However, research comparing historical and modern vital rates is needed to identify demographic factors that may explain declines. Based on matrix models, adult female survival has the greatest effect on the intrinsic rate of growth. Thus, we hypothesized that changes in annual female survival may explain contemporary declines in populations if that vital rate has decreased concomitant with population declines. We searched peer-reviewed literature, student theses and dissertations, and government reports for empirical estimates of annual female survival across the range of wild turkeys. Of the 51 resulting estimates of annual female survival across all subspecies, most were ≥0.6 through 2012, but 8 out of 10 estimates post-2012 were lower, consistent with observed trends in population trajectories. We used matrix models to illustrate that the observed change in annual female survival estimates indeed predicts a declining population trajectory, even when other important vital rates are at the high end of the published range of process variation. If the published annual female survival estimates represent those experienced more broadly in populations, our meta-analysis and matrix models indicate that decreases in this vital rate help explain declines observed in populations in some regions. The causative factor(s) associated with the apparent declines are unknown, and little research has evaluated management strategies to increase annual female survival. Accordingly, we encourage wild turkey researchers to focus efforts on producing robust estimates of annual female survival, cause-specific mortality, and rigorous evaluation of management strategies to improve annual female survival.

---

Wild turkey (*Meleagris gallopavo*) populations have recently declined throughout their range (Casalena et al. 2015; Eriksen et al. 2015, Chamberlain et al. 2022). For example, Londe et al. (2023) estimated a 9% annual population decline in the eastern subspecies (*Meleagris gallopavo silvestris*) based on a matrix model populated with vital rates published during 1970–2021. Such declines have catalyzed a widespread increase in wild turkey research and prompted several state agencies to adjust turkey hunting seasons and test hypotheses to explain observed declines (Quehl et al. 2024). Although the causative mechanisms explaining population declines remain unclear, hypotheses include changes in predator context, habitat, climate, disease, and hunter harvest (Casalena et al. 2015).

For wild turkeys, vital rates associated with the female segment of the population typically have the greatest influence on population trajectory. Accordingly, female-only matrix models often include estimates of adult and yearling survival, nesting rate, nest survival, renesting rate, clutch size, hatchability, and poult survival (Rumble et al. 2003, Pollentier et al. 2014, Londe et al. 2023). Several U.S. states have reported low poult-per-hen ratios, suggesting poor productivity may be negatively influencing populations (Byrne et al. 2015). Although nest and poult survival can influence population growth rates (Roberts et al. 1995, Roberts and Porter 1998, Pollentier et al. 2014), adult female survival commonly has the greatest effect on the intrinsic rate of growth (Roberts & Porter 1996, Rolley et al. 1998, Pollentier et al. 2014, Lehman et al. 2022, Londe et al. 2023). As such, population growth rates of wild turkeys are sensitive to even small changes in adult female survival (Suchy et al. 1983, Wakeling 1991, Vangilder and Kurzejeski 1995, Alpizar-Jara et al. 2001, Tyl et al. 2019, Londe et al. 2023).

Despite these findings, we are not aware of any attempts to determine if annual female survival has changed over time. Additionally, we are unaware of any attempts to determine whether decreases in annual female survival over time could explain observed declines in wild turkey populations. Thus, we performed a comprehensive review of the literature to compile all published estimates of adult female annual survival. We hypothesized that annual female survival has declined over time, particularly during the past two decades, mirroring the pattern observed in population declines. To explore how any observed changes in annual female survival estimates might affect population trajectory, we also populated female-only, stage-based matrix projection models to estimate the asymptotic population growth rate (λ), crossing low, average, and high annual female survival estimates with low, average, and high reproduction rates previously quantified by Londe et al. (2023).

## STUDY AREA

We used published data from the literature across the distribution of eastern, Rio Grande (*M. g. intermedia*), and Merriam’s (*M. g. merriami*) wild turkey subspecies, including data from 27 U.S. states and Canadian provinces (Figure 1). Data collection ranged from 1978–2023. Latitude ranged from 28.0204645°N–47.9535396°N, and site elevation ranged from 15–2,200m above sea level. Climate, vegetation, landownership, and landscape composition varied widely across the study area.

**Figure 1.**
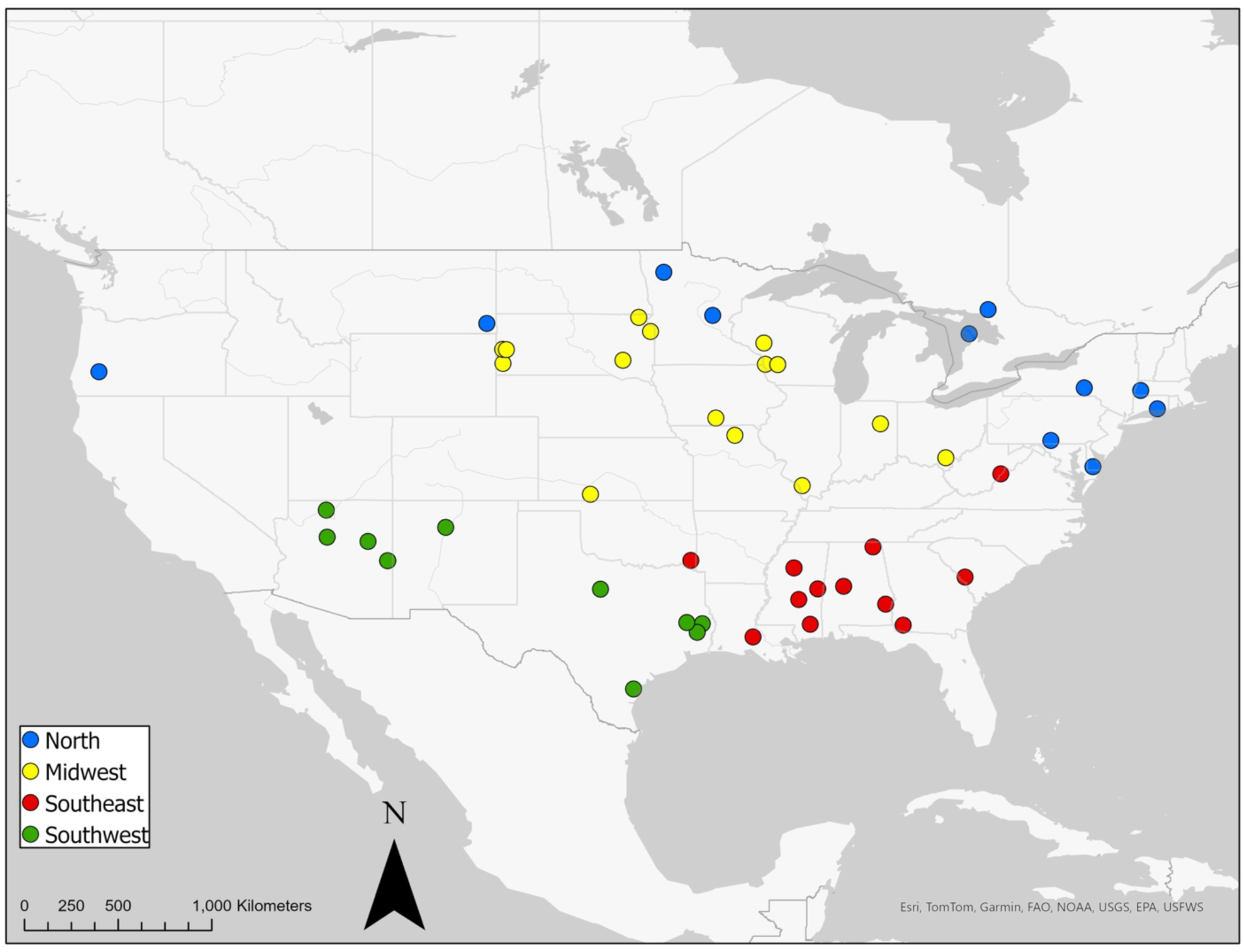
Location of studies that reported annual survival of female wild turkeys (*Meleagris gallopavo*) by region from 1978-2023.

## METHODS

### Literature review

We compiled literature in three ways. First, we downloaded the supplementary file provided by Londe et al. (2023) in their recent review of wild turkey vital rates from 1970 – 2021. Second, we conducted a literature search in Web of Science using the keywords “meleagris” and “survival.” Third, we downloaded all Wild Turkey Symposia Proceedings through May 2024 and searched for all manuscripts that may have included female survival estimates. In all studies including female survival estimates, we reviewed the literature cited for additional sources not found through other search methods. We screened over 2,000 manuscripts, theses, dissertations, and final reports for results related to annual or seasonal adult female survival. We only included studies estimating female survival from Very High Frequency (VHF) or Global Positioning System (GPS) tags, given their increased certainty relative to banding studies (Wightman et al. 2024).

### Data analysis

To assess trends in female survival over time, we used a meta-regression framework. When the authors provided an estimate of annual adult female survival and a measure of error (standard error, variance, confidence interval, or standard deviation + sample size), we pooled over the entire study duration to produce a single estimate of annual female survival. When the authors provided annual female survival estimates and sample sizes, we extracted annual female survival estimates and sample size for each year in the study. When the authors provided annual survival for each year of the study along with the total sample size, we assumed that samples were evenly spread across study years. When the authors only provided seasonal survival estimates, we estimated annual survival by calculating the product of all seasonal survival estimates. Due to varying approaches for reporting error across studies, we calculated the variance for all annual female survival estimates, which were proportions, as:

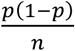

where *p* = annual female survival and *n* = sample size. To compare studies that reported multiple years of annual female survival with studies that only reported pooled values, we calculated the pooled annual survival and variance for all individual year-reporting studies using inverse-variance weighting (Mengersen et al. 2013). For all studies, we assigned pooled survival estimates to the final year of the study (Study Year). We also recorded the state or province where each study was conducted, which we collapsed into four mutually exclusive regions: North, Midwest, Southeast, and Southwest (Figure 1). We performed analyses using the subspecies in place of the region variable for all analyses and it did not affect the inferences (Figures S1 and S2); thus, we decided to group based on regions given that multiple subspecies often occur in the same state.

We conducted a meta-regression using annual survival as the response variable and Study Year and Region as the predictor variables. We weighted all regression models using the inverse variance from each study, which we normalized by dividing by the mean inverse variance of all studies (Mengersen et al. 2013). We used a beta distribution (Douma and Weedon 2019) because annual survival estimates were proportions (i.e., between 0 and 1).

We first tested for regional differences in female survival by fitting a model using only Region as a predictor variable. If we detected significant differences in regional survival using a Wald test, we then included Region as a predictor variable in subsequent models as both an additive and interactive term. If we failed to detect a difference in regional survival, we used Study Year as the only predictor variable in our models. Recognizing that non-linear patterns are common in ecology, we fit models with Study Year as a linear term and as an additive second- and third-order polynomial (Guthery and Bingham 2007). We ranked all models by Akaike Information Criteria for small sample sizes (AICc) and considered those with ΔAICc < 2.0 as competitive models (Burnham et al. 2011). We fit all models using package glmmTMB (Brooks et al. 2017) in Program R v4.4.1 (R Core Team 2024).

### Matrix models

After testing for a shift in adult female survival over time, we wanted to quantify the effect of differential adult female survival on population growth. Visually, it appeared female survival estimates in the literature began to decline around 2012 based on our meta-analysis. Thus, we created matrix models based on early (1978–2011) and late (2012–2024) estimates of female survival. We specifically used two matrix model approaches: deterministic and stochastic. We used the 2-stage female-based, prebirth pulse, Lefkovitch matrix model from Londe et al. (2023) as the basis for both approaches. First, we used a deterministic matrix model to compare effects of survival while reproduction was held constant. We calculated asymptotic growth rate (λ) for every pairwise combination of low, medium, and high survival and reproduction (0.25, 0.5, and 0.75 quantiles, respectively). The survival quantiles were calculated from all published survival rates during each period (early and late), regardless of subspecies and without consideration of how previous authors separated or combined age classes of females. The reproduction quantiles were taken from the process distribution published by Londe et al. (2023), which incorporated published data from 1970–2021 for adults of the eastern subspecies. Second, we used a stochastic matrix model approach to assess population growth potential under the full process distributions of survival and reproduction. For each of 10,000 iterations, we selected values of survival and reproduction from their process distributions, using normal distributions truncated at 0 and 1. Across the 10,000 iterations, we quantified the proportion of model runs that projected a stable or increasing population (i.e., λ ≥ 1).

## RESULTS

We identified 46 publications that reported hen survival, which resulted in 51 estimates of hen survival, spanning the years 1978– 2023 (S1). 25 of these publications were included in the Londe et al. (2023) review, and the remaining 21 were added via our literature review. Sixteen estimates (31%) came from the Midwest, 10 (20%) from the North, 14 (28%) from the Southeast, and 11 (22%) from the Southwest.

Pooled estimates of annual hen survival ranged from 21.9–85.8%, and the mean inverse-variance weighted annual survival across all studies was 61.1% (SE = 0.6%). In the first round of model selection, we detected significant differences in survival by Region (χ^2^ = 11.57, df = 3, p = .009). Therefore, in the next tier of model development, we included Region as both an additive and interactive effect with Study Year. Our top model indicated that the change in annual survival over time was non-linear (Table 1). The top model included Region as an additive fixed effect and showed that annual hen survival was 41.2% lower (β = -0.53, SE = 0.16, *p* < 0.001) in the North compared to the Midwest, which was our reference region. Annual hen survival in the Southeast and Southwest were similar to the Midwest (*p* = 0.25, *p* = 0.36, respectively). The top model also included Study Year (β = -1.12, SE = 0.45, *p* = 0.01), Study Year^2^ (β = -0.53, SE = 0.49, *p* = 0.28), and Study Year^3^ (β = -1.22, SE = 0.47, *p* = 0.009; Figure 2).

**Figure 2.**
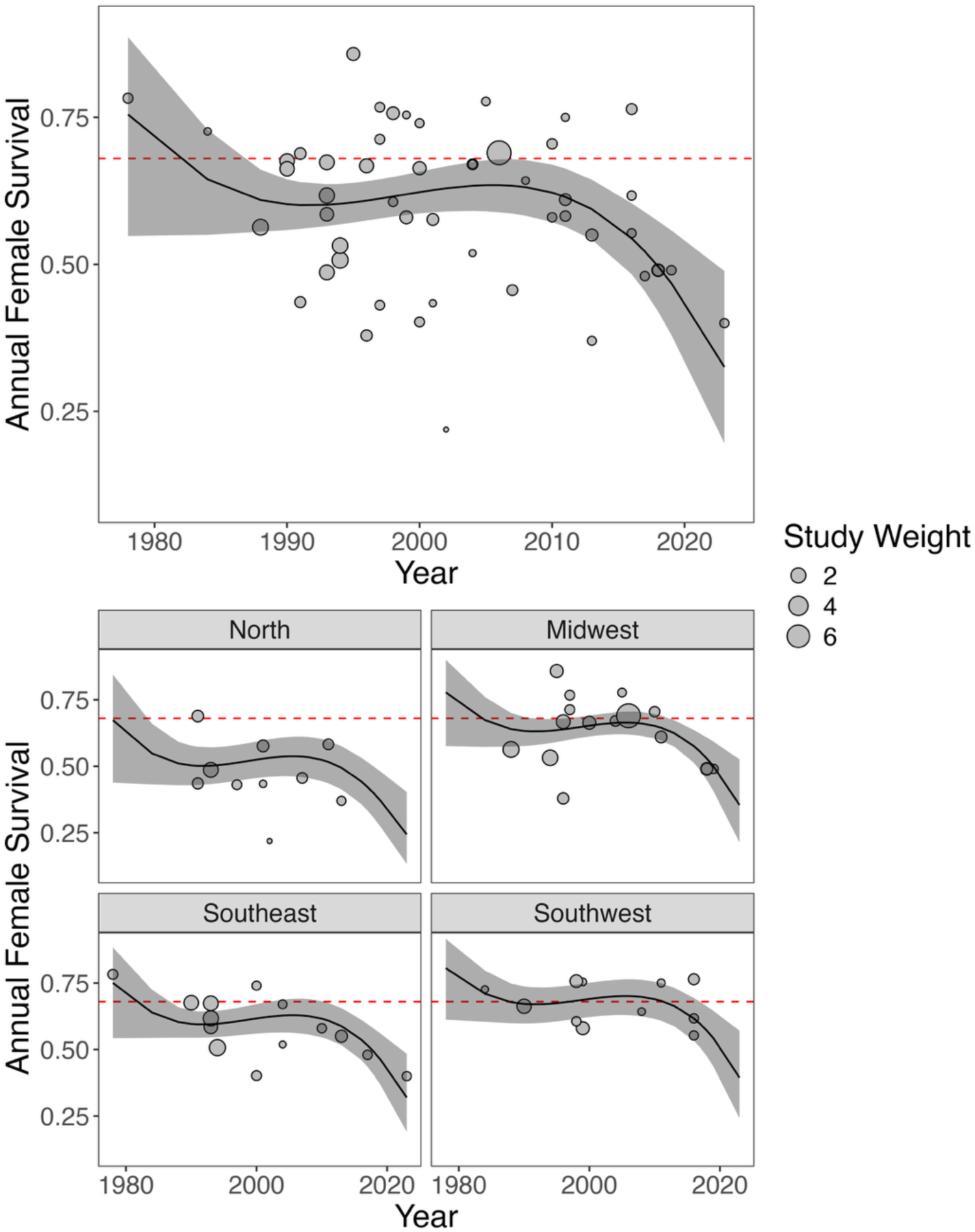
Model results from meta-regression showing estimated marginal means of annual female survival from 1978 to 2023. The top figure illustrates the estimated marginal means of annual female survival across all studies, while the lower figures display the estimated marginal means of annual female survival across four different regions of North America. The dashed red line (0.68) indicates Lehman et al.’s (2022) estimate of the minimum female survival rate required to maintain a stable turkey population. The point size in the figures reflects the weight of each study in the meta-regression, calculated as the standardized inverse variance.

**Table 1.** Model selection table providing the number of parameters (K), log-likelihood (logLik), Akaike’s Information Criteria for small sample sizes (AICc), ΔAICc, and model weight (w) for beta regression models investigating the impacts of time and region and annual female turkey survival in North America. Models are ranked by AIC and only models with ΔAICc < 2 were considered top models.

| Model | K | logLik | AICc | ΔAICc | w |
| --- | --- | --- | --- | --- | --- |
| Survival ~ Year + Year<sup>2</sup> + Year<sup>3</sup> + Region | 8 | 52.666 | -85.9 | 0 | 0.719 |
| Survival ~ Year + Region | 6 | 48.51 | -83.1 | 2.79 | 0.178 |
| Survival ~ Year + Year<sup>2</sup> + Region | 7 | 49.195 | -81.8 | 4.12 | 0.092 |
| Survival ~ Year * Region | 9 | 49.583 | -76.8 | 9.13 | 0.007 |
| Survival ~ Year * Region + Year<sup>2</sup> * Region | 13 | 55.588 | -75.3 | 10.57 | 0.004 |
| Survival ~ Year * Region + Year<sup>2</sup> * Region + Year<sup>3</sup> * Region | 17 | 56.805 | -61.1 | 24.84 | 0 |

### Matrix models

Using the 0.25, 0.5, and 0.75 quantiles (low, medium, and high, respectively) of survival from each of the early and late periods (Table 2) for our deterministic matrix model approach, we determined that only survival from the early period was sufficient to support a stable or growing population (i.e., λ ≥ 1; Figure 3), and this was only when survival was matched with the high reproduction rate. By contrast, no value of survival in the late period could support a stable or growing population, even when combined with high reproduction. When holding survival at its median value for the early period, recruitment needed to be in the 72nd percentile to maintain a stable population (i.e., λ = 1), and the 93rd percentile for the late period to achieve a stable population. When comparing λ between the early and late periods for each combination of low, medium, and high reproduction and survival, λ decreased 9–16% over time (Figure 3). Using the stochastic matrix model approach, we determined that mean λ decreased from 0.93 to 0.79 between the early and late periods (Figure 4). Additionally, 75% of model runs in the early period showed a decreasing population (λ < 1), and 96% of runs in the late period showed a decreasing population (Figure 4).

**Figure 3.**
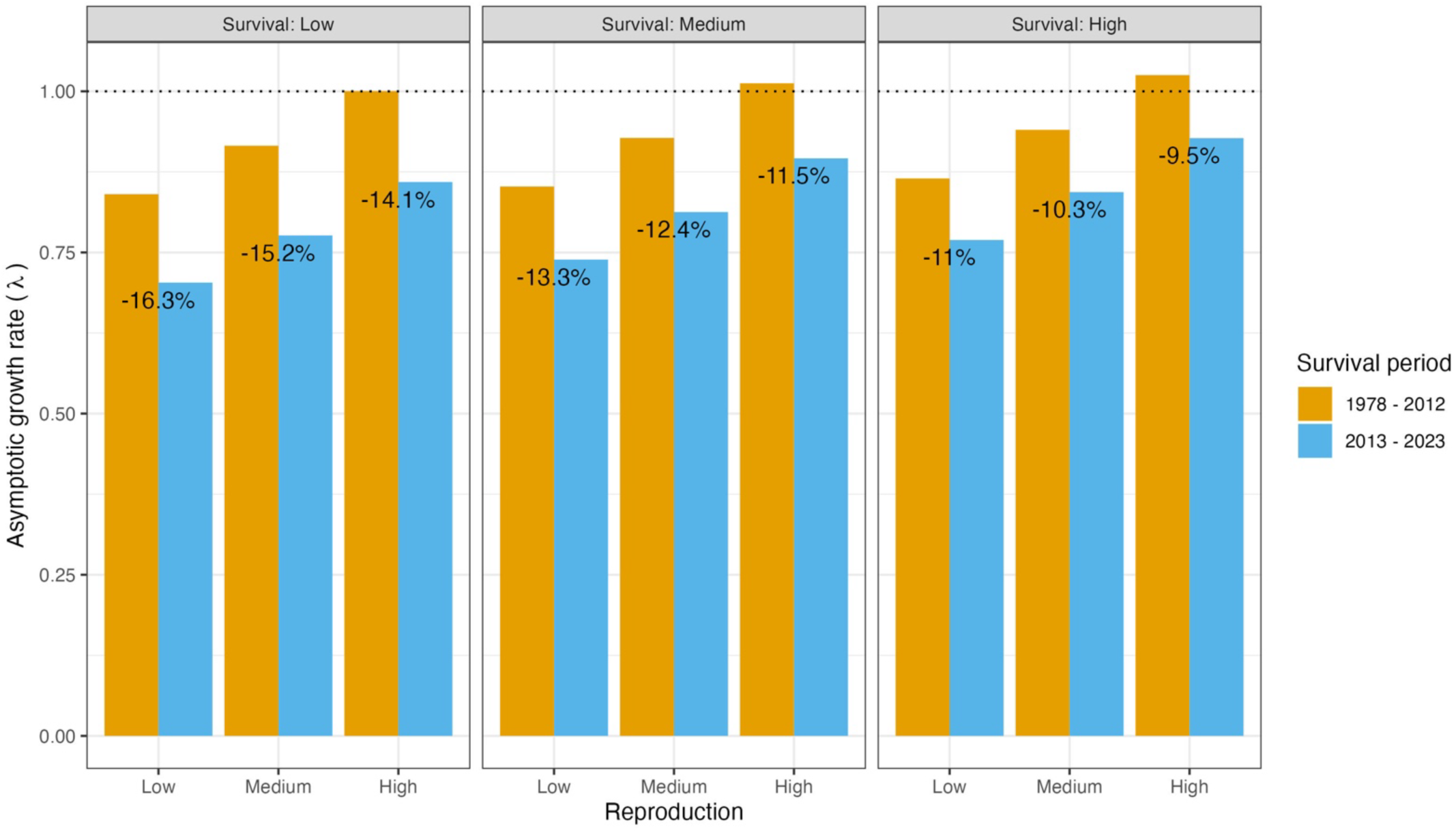
Asymptotic growth rate (λ) from the deterministic matrix model under low, medium, and high survival rates (0.25, 0.5, and 0.75 quantiles, respectively) from the early period (1978–2012) and late period (2013–2023), matched with low, medium, and high reproduction rates (0.25, 0.5, and 0.75 quantiles, respectively) from 1970–2021 (from Londe et al. [2023]). Percentages shown on the bars are the percent decrease in λ between the early (orange) and late (blue) periods for each respective pairing.

**Figure 4.**
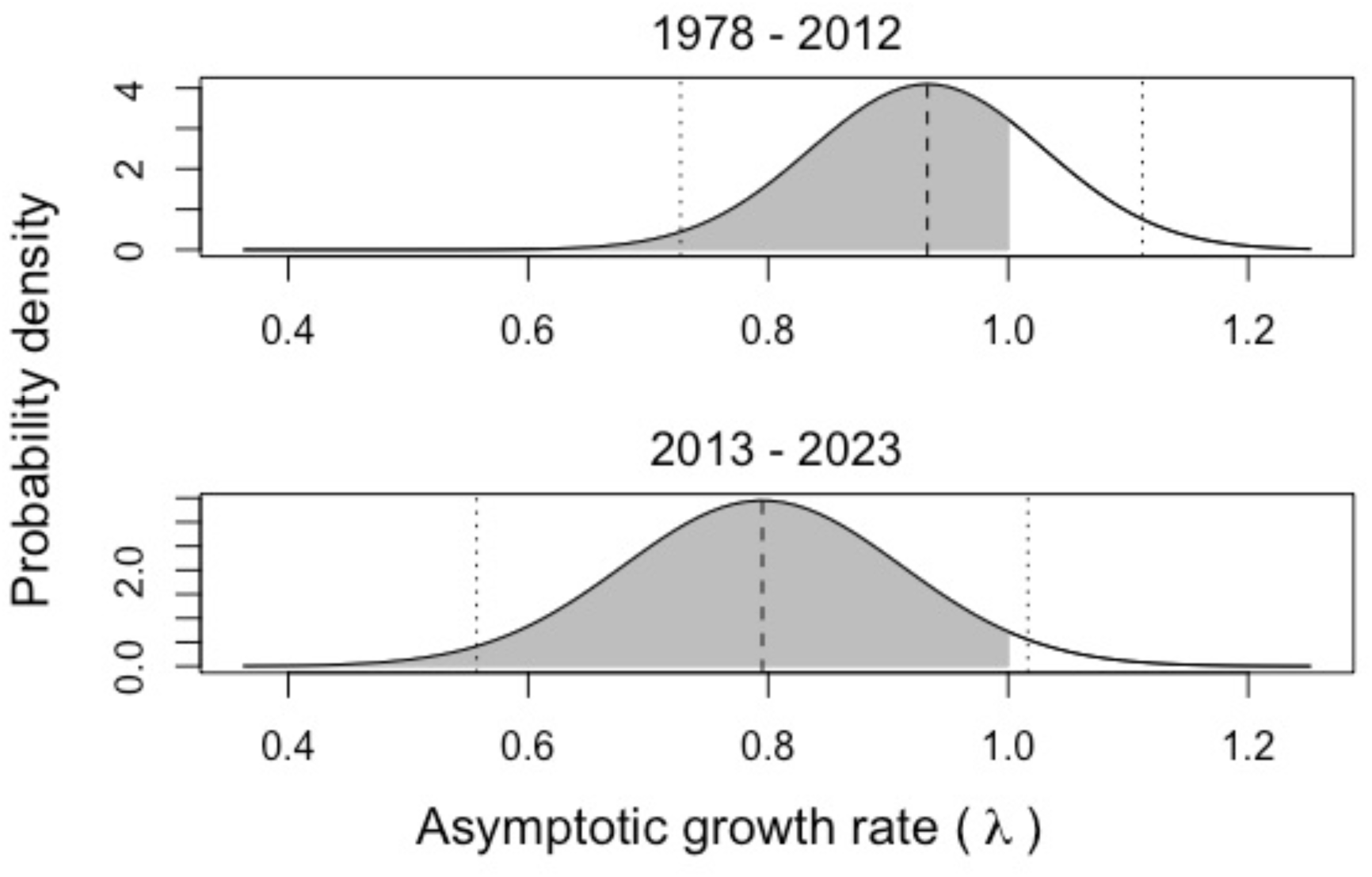
Estimated asymptotic growth rate (λ) for wild turkey using survival from two time periods: 1978–2012 and 2013–2023. Dashed lines represent mean λ from 10,000 iterations of the stochastic matrix model (0.93 for the early period and 0.79 for the late period); dotted lines represent 95% confidence intervals. Shaded gray areas indicate values representing a declining population (λ < 1), and unshaded area shows area indicating an increasing population (λ > 1). In the early period, 75% of model runs indicated a declining population, and in the late period 96% of model runs indicated a declining population.

**Table 2.** Annual survival of adult female wild turkey by quantile (as well as mean and standard deviation [SD]) for two time periods (1978–2012 and 2013–2023).

| Period | Quantile | Annual Survival |
| --- | --- | --- |
| 1978–2012 | 0.25 | 0.639 |
|  | 0.5 | 0.650 |
|  | 0.75 | 0.662 |
|  | Mean (SD) | 0.655 (0.030) |
| 2013–2023 | 0.25 | 0.507 |
|  | 0.5 | 0.541 |
|  | 0.75 | 0.570 |
|  | Mean (SD) | 0.523 (0.093) |

## DISCUSSION

Our results indicated that temporal trends in female survival help explain wild turkey population declines. The change in hen survival over time is interesting and has implications for population performance. Adult female survival is the most important factor influencing wild turkey population growth rates (Londe et al. 2023), and small changes to this vital rate strongly influence λ. Indeed, our matrix models confirmed that the observed changes in published annual hen survival estimates would correspond with a declining population growth rate, even when other important vital rates are at the upper end of their published range of the process distribution. As noted by Londe et al. (2023), it is important to consider that the variation in vital rates that constitute reproductive performance cannot be ignored. Though turkeys are shorter lived than most ungulates, demographic research in relatively long-lived species has shown that greater variation inherent in some vital rates (e.g., fawn/calf survival) gives them a disproportionate effect on population trajectory (Gaillard et al. 1998, 2000, Raithel et al. 2007, Chitwood et al. 2015), especially when it comes to management. The natural variability in vital rates is the primary determinant of spatial and temporal variation in λ (Raithel et al. 2007). Therefore, management actions should be more effective at moving a vital rate when it has more variation with which to work. Continued research into how management can improve turkey reproduction (the more variable rate) will be critical, but adult females must survive for reproduction to matter.

Several aspects of our meta-analysis give us confidence that the studies included represent state- and regional-level population trends. First, the declines in survival we observed over time generally matched reported declines in time and space (Casalena et al. 2015, Eriksen et al. 2015, Chamberlain et al. 2022). For example, Casalena et al. 2015 found that turkey harvest rates (a proxy for turkey abundance), declined by 25% in Mid-Atlantic states since the early 2000s. Additionally, among states that consistently reported turkey abundance, there was an estimated 3% decline in population size between 2014 and 2019 (Chamberlain et al. 2022). Second, in most cases where multiple reports from the same state were available, survival was greater in the earlier study (Figure 5). We only found one instance where an historical and modern female survival estimate was reported from the same study site. In that South Dakota population, annual adult female survival decreased from 0.67 during 1999–2001 (Shields and Flake 2006) to 0.49 during 2017–2019 (Tyl et al. 2023). Thus, we believe the negative temporal trend in adult female survival we observed is likely representative of real demographic change over time.

**FIGURE 5:**
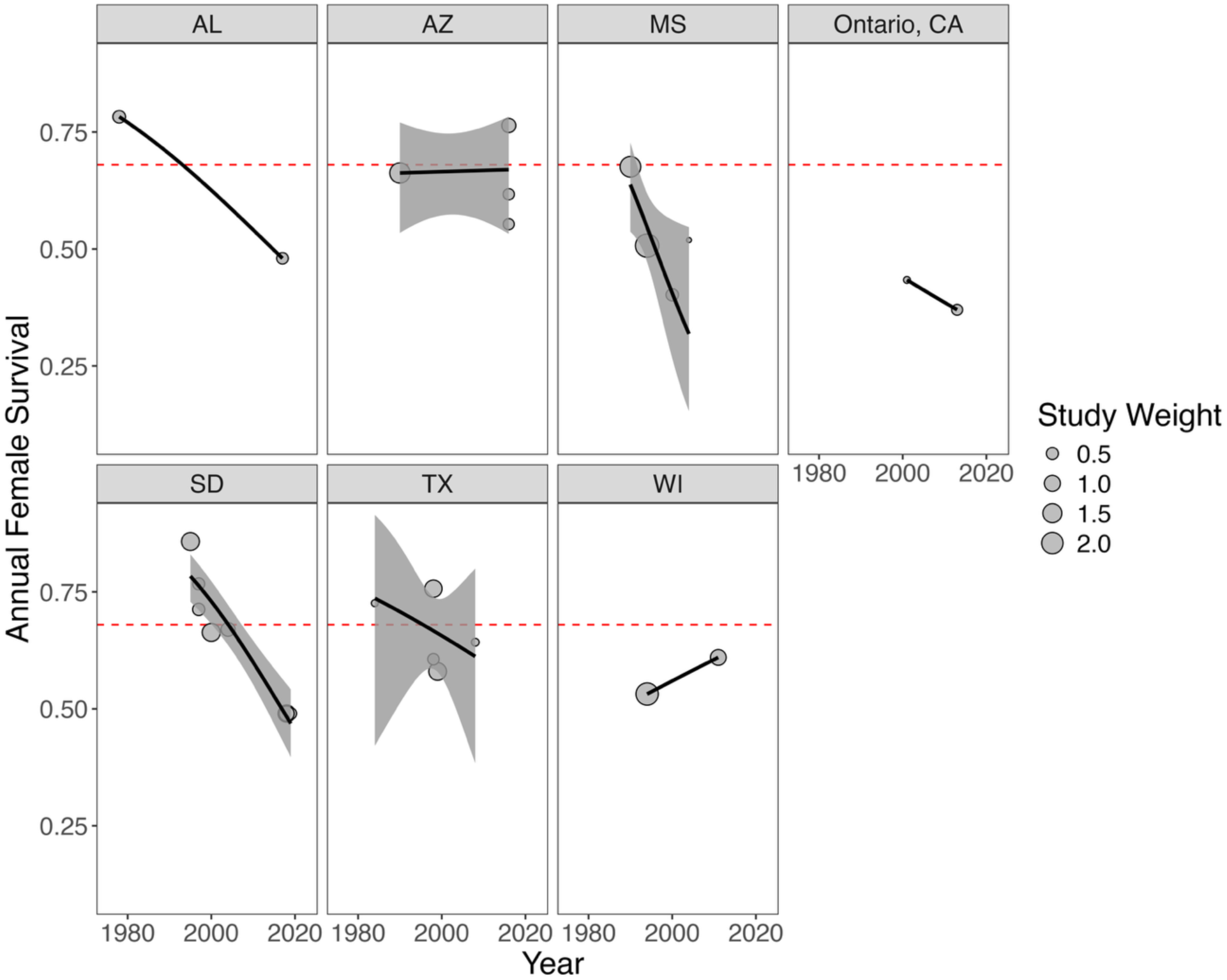
Model results from meta-regression showing annual female survival estimates from 1978–2023 for North American states and provinces with repeated temporal measures that were at least 10 years apart. The dashed red line (0.68) shows Lehman et al.’s (2022) estimate of minimum female survival to maintain a stable turkey population. Point size indicates each study’s weight in the meta-regression, which was calculated as the standardized inverse variance.

Historically, managers have manipulated female survival in wild turkey populations by adjusting hunter harvest (Vangilder and Kurzejeski 1995). Conservative fall harvest rates of ≤10% have been recommended to maintain population growth; however, these harvest rates were based on historical female survival rates ≥56% (Vangilder and Kurzejeski 1995, Alpizar-Jara et al. 2001, McGhee et al. 2008). In recognition of the role of hunter harvest in female survival, combined with declining populations, many states currently allow limited or no female harvest. Thus, most managers lack the flexibility to increase female survival through regulatory mechanisms. Given the declines in female survival in the literature, we agree with Londe et al. (2023) that female harvests may not be sustainable in areas where turkey populations are declining.

Mechanisms to explain recent declines in female survival are unknown, but we believe several factors are plausible. Negative density dependence may contribute to population declines, but Byrne et al. (2015) reported increasing female survival independent of population size from 1981–2012 in the southeastern U.S., indicating the temporally decreasing female survival we observed may not be related to density. Interestingly, Byrne et al. (2015) included only two estimates from studies at the start of the period where we documented a decline in female survival, and it appears the negative trend became evident after their time series ended. Food availability could influence adult survival, and there have been well-documented declines in both insect and insectivorous bird populations; the spatial and temporal extent of insect declines align with declines in wild turkey populations (Van Klink et al. 2020, Wagner 2020, Tallamy and Shriver 2021). Furthermore, acorns are an important fall and winter food resource for turkeys (Meanley 1956, Hurst 1992), and overstory oaks (*Quercus* spp.) have been declining in many areas (McShea et al. 2007, Haavik et al. 2015, Alexander et al. 2021). It also is possible that baiting and supplemental feeding, which only recently became widespread practice in many states (Arkansas Game and Fish Commission 2019, South Carolina Department of Natural Resources 2020, Huang et al. 2022), could negatively affect female survival. Feeding may expose females to greater predation risk during nesting (Cooper and Ginnett 2000), while also posing a health threat by increasing disease and toxin exposure (Huang et al. 2022; Huang et al. 2021). There has also been increased concern over several diseases that could negatively affect female survival (MacDonald et al. 2022); however, little information is available to understand how exposure to diseases has changed over space and time.

Changes in predation risk associated with changes in predator abundance or communities is also a plausible mechanism for the trend we observed. Female survival is generally lowest during the spring reproductive period (Vander Haegen et al. 1988, Yarnall et al. 2020, Tyl et al. 2023), and multiple studies point to predators as the primary source of cause-specific mortality (Kurzejeski et al. 1987, Miller et al. 1998, Hubbard et al. 1999, Nguyen et al. 2003, Moore et al. 2010). There are examples of a changing wild turkey predator context across our study period. For example, coyotes (*Canis latrans*) are a common predator of females (Miller et al. 1995, Delahunt 2011, Pollentier et al. 2014), the timeline of their range expansion into the eastern U.S. coincides with turkey population declines in some areas (Hody and Kays 2018), and coyote abundance was negatively correlated with wild turkey abundance in Mississippi (Wang et al. 2023). By contrast, coyote range expansion seems insufficient to explain turkey survival decreases range-wide, as recent changes to female survival also have occurred within the historical range of coyotes (e.g., Shields and Flake 2006, Tyl et al. 2023).

Based on recent reductions in landscape coverage of herbaceous plant communities (Keyser et al. 2019, Bernath-Plaisted et al. 2023), along with the nesting cover they provide (Crawford et al. 2021), another hypothesis is that predation on adult female turkeys may be increasing in response to recent reductions in landscape coverage of nesting and brooding cover. Indeed, occupancy probability for mammalian turkey predators was decreased in burned areas as compared to non-burned areas in a recent study (Boone et al. 2024). Climate also may influence survival, as increased precipitation can increase female and nest predation, perhaps because mammalian predators are more effective at locating nesting females via olfaction during or immediately following rain events (Roberts et al. 1995, Lehman et al. 2008, Webb et al. 2012, Yarnall et al. 2020). Overall, it is likely that several of the proposed mechanisms outlined above have interactive or additive effects that contributed to the temporal decline in annual adult female survival we observed. Specifically, reduced food availability and disease could decrease female body condition, survival, and ability to escape predators, while changes in baiting and feeding practices, habitat, and climate have simultaneously increased predation risk. While other mechanisms may play a role, we suggest these hypotheses are worth exploring as mechanisms to explain declines in female survival.

One potential limitation of this meta-analysis approach is that the studied populations may not be representative of the broader trends in unstudied populations. For example, much of the research conducted on wild turkeys in the past was associated with recently restocked populations that were stable or increasing (Vander Haegen et al. 1988, Roberts et al. 1995, Shields et al. 2006), whereas more contemporary work may have focused on declining populations or those perceived as declining by biologists (Zenas 2018, Lehman et al. 2022, De Filippo 2024). Methods have also been inconsistent, as several studies only provided survival estimates for a combination of adult and juvenile females (Vangilder et al. 1995, Lehman et al. 2000, Nguyen et al. 2003), whereas more recent studies commonly separate analyses by age class (Pollentier et al. 2014, Little et al. 2016, Tyl et al. 2023). Many studies included in our analyses reported no difference in adult and juvenile female survival (Hennen and Lutz 2000, Kane et al. 2007, Delahunt 2011, Yarnall et al. 2020, Tyl et al. 2023), whereas others did (Vander Haegen et al. 1988). To ensure this was not an issue in our approach, we also ran the analyses excluding studies that combined age groups; the resulting patterns over time mirrored the analysis of the full dataset (Figure S3). Changes in technology also may have influenced our results, as earlier studies primarily used VHF transmitters, whereas several modern studies used both GPS and VHF transmitters (Niedzielski and Bowman 2014, Zenas 2018, De Filippo 2024). Nonetheless, VHF and GPS data should provide similar survival estimates (Latham et al. 2015), especially given that studies employing GPS transmitters often rely on a VHF beacon for mortality checks (Niedzielski and Bowman 2014). Moreover, we excluded banding studies in favor of radiotag studies as a precautionary measure to avoid biases associated with monitoring methods.

Most previous research and management to increase turkey populations focused on improving productivity, but our results indicate focusing efforts on practices that increase female survival may be just as important. Pollentier et al. (2014) suggested that continued restoration and enhancement of high-quality habitat with early successional vegetation has the greatest potential to positively influence population growth by improving nest and poult survival. Such improvements to nesting and brood-rearing areas also could improve female survival by reducing mortality risks while incubating eggs and raising poults (Pollentier et al. 2014), as greater female predation rates may occur during nesting if cover is limited (Hohensee 1998). Similar habitat manipulations were suggested for Merriam’s turkeys in the West, and vegetation treatments directed at improving body condition of females through improved abundance and distribution of food would likely increase nesting rates (Wakeling 1991, Wakeling and Rodgers 1995, Hoffman et al.1996, Rumble et al. 2003). However, there is limited information on whether vegetation influences female survival, especially during incubation. Lohr et al. (2020) reported no effect of vegetation height on survival of incubating females, but multiple vegetation covariates such as density and visual obstruction may interact to influence survival (Fuller et al. 2013). Even if vegetation structure and composition has limited influence on nest survival (Keever et al. 2023), it could have a large impact on population growth if vegetation conditions reduce hen predation rates during nesting. Importantly, there have been no manipulative studies to determine the effect size of any management strategies on female survival, other than those associated with hunter harvests.

## MANAGEMENT IMPLICATIONS

Our results suggest a couple of opportunities for future research and management of wild turkeys. First, female survival is a critical metric influencing wild turkey population growth, which means that any management actions, harvest-related or otherwise, that improve female survival will have positive effects on population performance. Restricting or removing harvest of females (bearded or not) from spring and fall seasons is an easy first step in areas where female harvest is currently legal and declines are apparent. Second, mechanistic understanding about how management can move the needle on reproductive vital rates is critical. High-end reproductive values can offset reduced hen survival but only if those values are sustained. Based on current data and anecdotal reports across the country, periodic booms in reproduction will not be sufficient to stall observed declines. Therefore, we recommend renewed focus on the landscape variables most likely to affect hen nutritional condition, nest success, and poult survival. Where possible, experimental design should be prioritized to help isolate variables affecting wild turkey vital rates. Here, we challenge the wild turkey research community to develop rigorous, manipulative studies and take advantage of natural experiments, when possible, to isolate causative factors linked to female survival and reproductive success and to evaluate the effectiveness of corrective management actions.

---

**Supplementary Table 1.** Average female wild turkey (*Meleagris gallopavo*) annual survival estimate, standard error (SE), final study year, U. S. state or Canadian province where study was conducted, assigned Region, and Subspecies (Subsp.) for studies from 1978–2023 used in the meta-analysis. Regions included midwest (MW), north (N), southeast (SE), and southwest (SW) and subspecies included eastern (E; *M. g. silvestris*), Rio Grande (R; *M. g. intermedia*), and Merriam’s (M; *M. g. merriami*) wild turkeys.

| Article | Survival | SE | Year | State/Province | Region | Subsp. |
| --- | --- | --- | --- | --- | --- | --- |
| Everett et al. 1980 | 0.78 | 0.061 | 1978 | AL | SE | E |
| Ransom et al. 1987 | 0.73 | 0.108 | 1984 | TX | SW | R |
| Vangilder and Kurzejeski 1995 | 0.56 | 0.029 | 1988 | MO | MW | E |
| Wakeling 1991 | 0.66 | 0.034 | 1990 | AZ | SW | M |
| Palmer et al. 1993 | 0.68 | 0.033 | 1990 | MS | SE | E |
| Thompson 1993 | 0.44 | 0.051 | 1991 | MT | N | M |
| Keegan and Crawford 1999 | 0.69 | 0.047 | 1991 | OR | N | R |
| Pack et al. 1999 | 0.59 | 0.037 | 1993 | WV | SE | E |
| Pack et al. 1999 | 0.62 | 0.031 | 1993 | VA | SE | E |
| Pack et al. 1999 | 0.67 | 0.033 | 1993 | WV | SE | E |
| Roberts et al. 1995 | 0.49 | 0.032 | 1993 | NY | N | E |
| Miller et al. 1998 | 0.51 | 0.029 | 1994 | MS | SE | E |
| Wright et al. 1996 | 0.53 | 0.030 | 1994 | WI | MW | E |
| Leif 2000 | 0.86 | 0.040 | 1995 | SD | MW | E |
| Hubbard et al. 1999 | 0.67 | 0.036 | 1996 | IA | MW | E |
| Hennen and Lutz 2000 | 0.38 | 0.050 | 1996 | KS | MW | R |
| Lehman et al. 2000 | 0.71 | 0.062 | 1997 | SD | MW | E |
| Lehman et al. 2000 | 0.77 | 0.064 | 1997 | SD | MW | R |
| Spohr et al. 2004 | 0.43 | 0.064 | 1997 | CT | N | E |
| Hohensee 1999 | 0.61 | 0.068 | 1998 | TX | SW | R |
| Kelly 2001 | 0.76 | 0.041 | 1998 | TX | SW | E |
| Feuerbacher et al. 2005 | 0.58 | 0.040 | 1999 | TX | SW | E |
| Whiting et al. 2005 | 0.75 | 0.102 | 1999 | TX | SW | E |
| Inglis 2001 | 0.40 | 0.062 | 2000 | MS | SE | E |
| Shields and Flake 2006 | 0.66 | 0.039 | 2000 | SD | MW | E |
| Moore 2006 | 0.74 | 0.073 | 2000 | SC | SE | E |
| Casalena et al. 2005 | 0.58 | 0.046 | 2001 | PA | N | E |
| Nguyen et al. 2003 | 0.43 | 0.109 | 2001 | ON, CA | N | E |
| Kane et al. 2007 | 0.22 | 0.169 | 2002 | MN | N | E |
| Lehman et al. 2005 | 0.67 | 0.054 | 2004 | SD | MW | M |
| Wilson et al. 2005 | 0.67 | 0.075 | 2004 | LA | SE | E |
| Holder 2006 | 0.52 | 0.118 | 2004 | MS | SE | E |
| Humberg et al. 2009 | 0.78 | 0.079 | 2005 | IN | MW | E |
| Reynolds and Swanson 2010 | 0.69 | 0.017 | 2006 | OH | MW | E |
| Parent et al. 2011 | 0.46 | 0.054 | 2007 | MN | N | E |
| Isabelle 2010 | 0.64 | 0.101 | 2008 | TX | SW | E |
| Delahunt 2011 | 0.71 | 0.057 | 2010 | IL | MW | E |
| Byrne and Chamberlain 2018 | 0.58 | 0.067 | 2010 | LA | SE | E |
| Ludwig 2012 | 0.58 | 0.055 | 2011 | DE | N | E |
| Peyton et al. 2014 | 0.75 | 0.092 | 2011 | NM | SW | M |
| Pollentier et al. 2014 | 0.61 | 0.046 | 2011 | WI | MW | E |
| Little et al. 2016 | 0.55 | 0.044 | 2013 | GA | SE | E |
| Niedzielski and Bowman 2014 | 0.37 | 0.072 | 2013 | ON, CA | N | E |
| McCall et al. 2020 | 0.55 | 0.071 | 2016 | AZ | SW | M |
| McCall et al. 2021 | 0.62 | 0.069 | 2016 | AZ | SW | M |
| McCall et al. 2022 | 0.76 | 0.053 | 2016 | AZ | SW | M |
| Zenas 2018 | 0.48 | 0.068 | 2017 | AL | SE | E |
| Lehman et al. 2022 | 0.49 | 0.041 | 2018 | SD | MW | M |
| Yarnall et al. 2020 | 0.49 | 0.048 | 2018 | SD | MW | M |
| Tyl et al. 2023 | 0.49 | 0.067 | 2019 | SD | MW | E |
| De Filippo 2024 | 0.40 | 0.071 | 2023 | OK | SE | E |

**Supplementary Figure 1.**
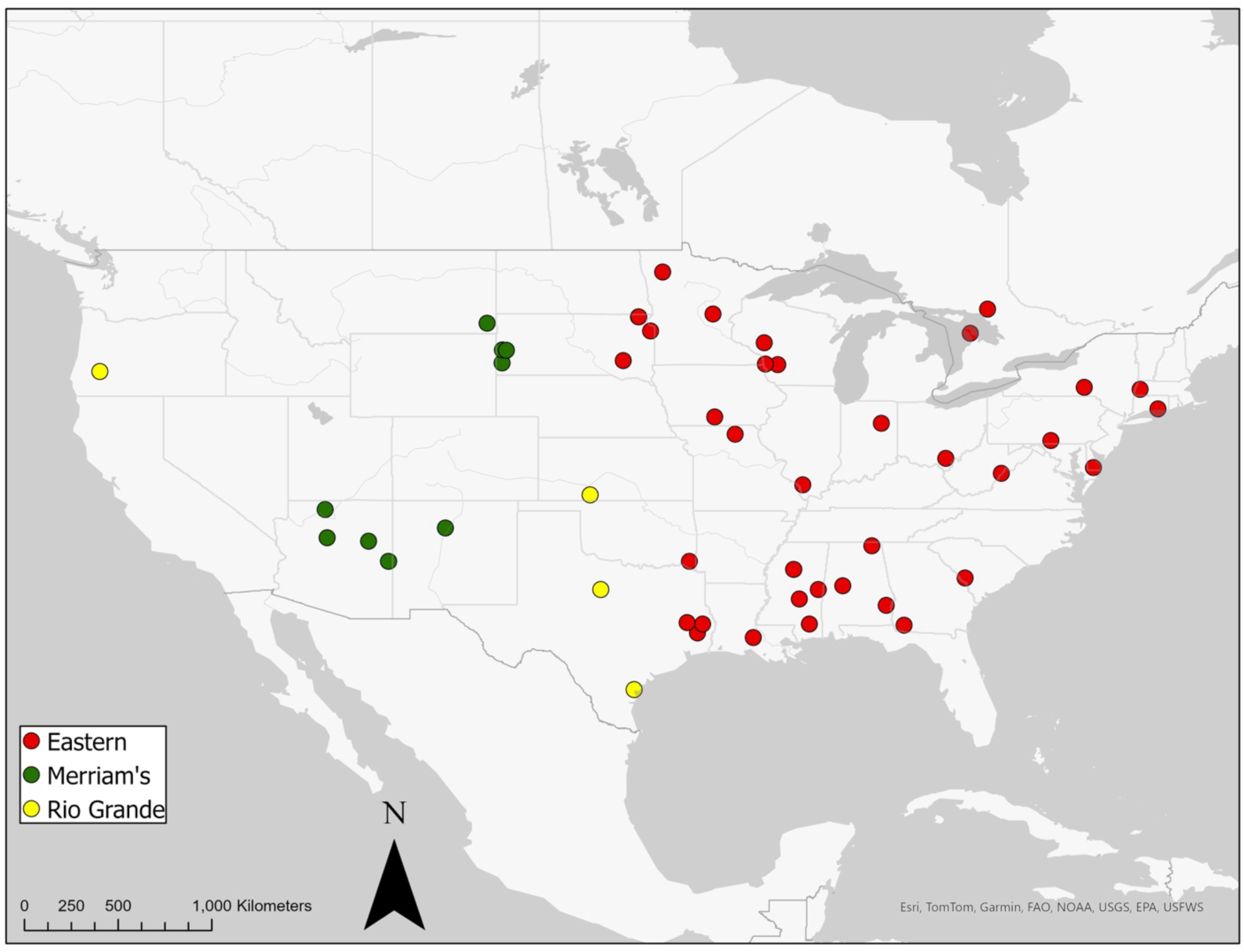
Location of studies that reported annual survival of female eastern wild turkey (*Meleagris gallopavo silvestris*), Merriam’s wild turkey (*M. g. merriami*), and Rio Grande wild turkey (*M. g. intermedia*).

**Supplementary Figure 2.**
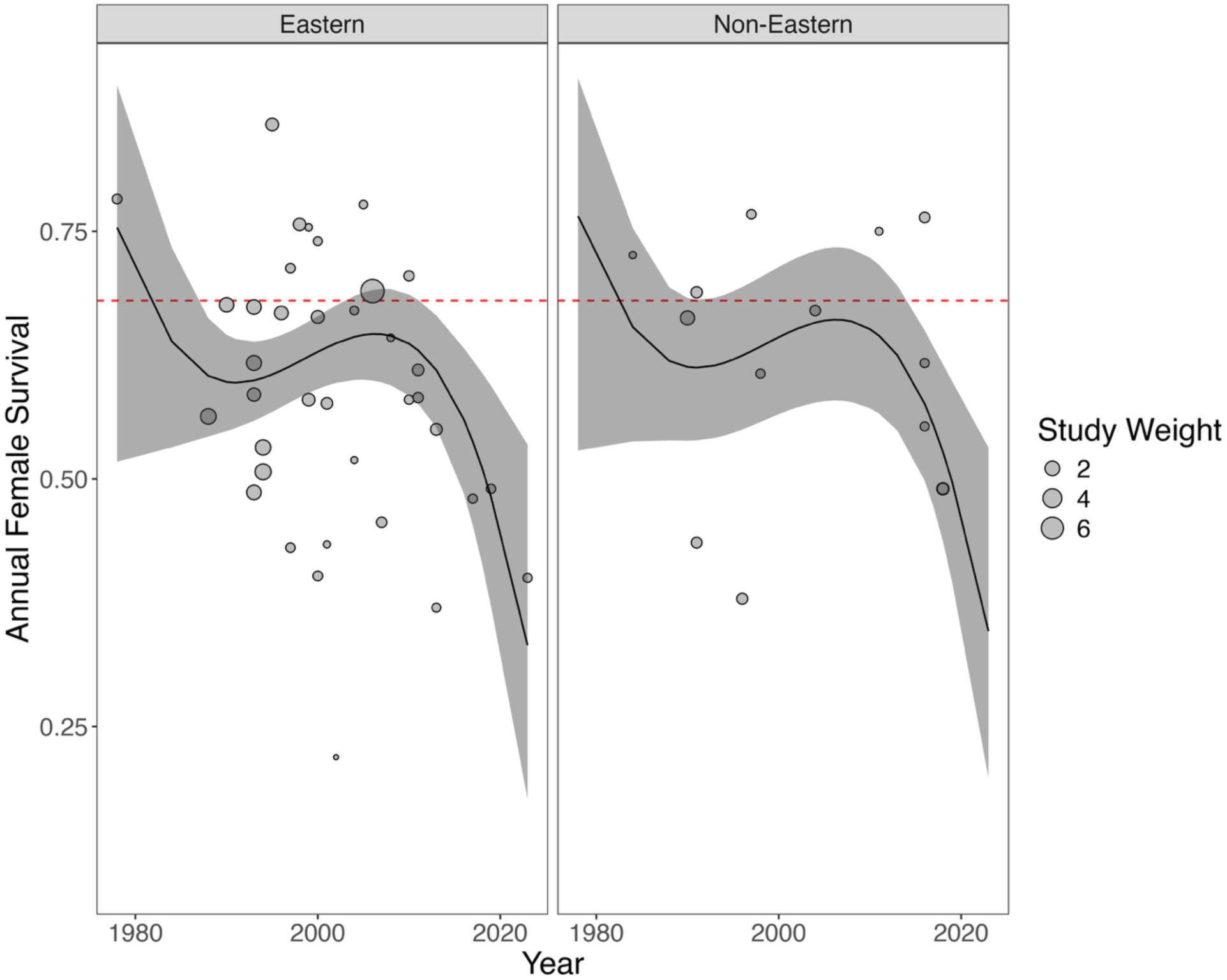
Model results from meta-regression showing estimated marginal means of annual survival from 1978 to 2023 for female eastern wild turkey (*Meleagris gallopavo silvestris*; left panel) and annual survival of female Merriam’s wild turkey (*M. g. merriami*) and Rio Grande wild turkey (*M. g. intermedia*) combined (right panel) from 1978-2023. Point size indicates each study’s weight in the meta-regression, which was calculated as the standardized inverse variance.

**Supplementary Figure 3.**
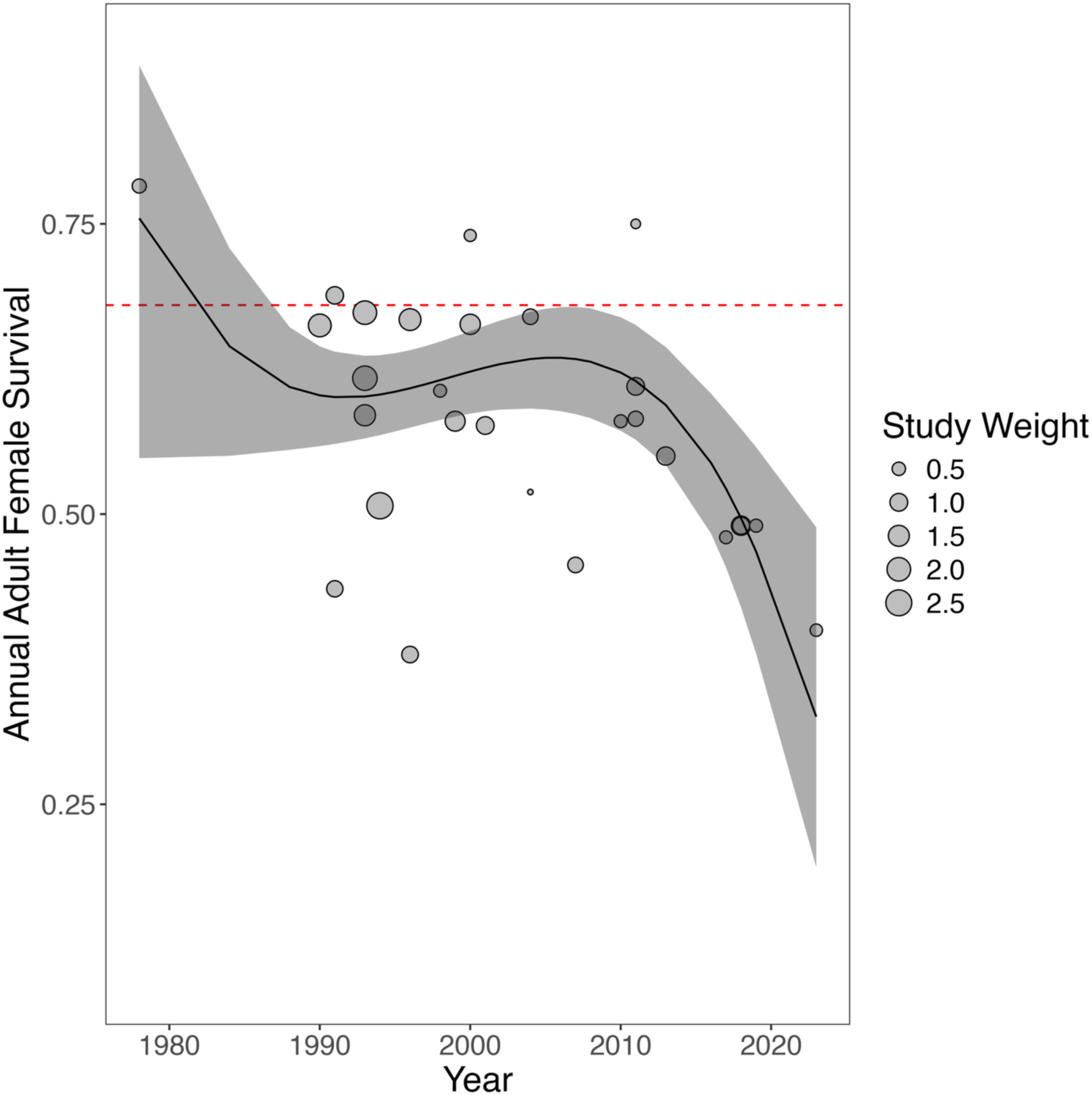
Model results from meta-regression showing annual female survival estimates for adults only from 1978–2023. The dashed red line (0.68) shows Lehman et al.’s (2022) estimate of minimum female survival to maintain a stable turkey population. Point size indicates each study’s weight in the meta-regression, which was calculated as the standardized inverse variance.

